# Spatial confinement induces oscillatory migration of epidermal keratinocytes and forms a three-compartment epithelial structure with epithelial–mesenchymal transition dynamics

**DOI:** 10.1101/2024.11.12.623158

**Authors:** Takuma Nohara, Sota Itamoto, Junichi Kumamoto, Yosuke Mai, Mayuna Shimano, Sora Kato, Hiroyuki Kitahata, Hideki Nakamura, Shota Takashima, Mika Watanabe, Masaharu Nagayama, Tsukasa Oikawa, Hideyuki Ujiie, Ken Natsuga

## Abstract

Epithelial cells undergo epithelial–mesenchymal transition (EMT) during migration and regain their epithelial phenotype in the post-migration phase (mesenchymal–epithelial transition; MET). We established an experimental system that reproduces a three-compartment epithelial structure comprising the original epithelium, its EMT state, and its MET state. Keratinocytes (KCs), skin epithelial cells, placed on a microporous membrane migrated through 3.0-µm or larger micropores. The 3.0-µm-pored membrane induced an epithelial structure with three distinct states: stratified KCs above the membrane, KCs showing EMT within the micropores, and a new stratified epithelium under the membrane. The membrane with larger micropores failed to maintain the three-compartment epithelial structure. Live imaging revealed that KCs moved in an oscillatory manner, with actin-rich filopodia-like structures extending into and out of the 3.0-µm micropores, while the cells migrated unidirectionally into larger micropores. Piezo1 and keratin 6 were identified as negative modulators of KC entry into and exit from the 3.0-µm micropores. These results demonstrate that non-cancerous epithelial cells migrate through confined spaces in an oscillatory manner, which might contribute to the formation of a three-compartment epithelial structure that recapitulates key aspects of wound healing.

## Introduction

Organ development and regeneration require cell migration to areas where other cells are absent, a phenomenon known as collective cell migration (Ilina and Friedl 2009). Epithelial cells typically undergo epithelial–mesenchymal transition (EMT) to facilitate this migration. Once the cells are in the appropriate region, they regain their epithelial phenotype through mesenchymal–epithelial transition (MET) (Leggett et al. 2021; Kalluri and Weinberg 2009; Dongre and Weinberg 2019). The epidermis, a stratified epithelium of the skin, serves as a barrier for living organisms (Natsuga 2014), and its major constituents are keratinocytes (KCs). KC stem cells located at the dermo-epidermal junction provide proliferating and differentiating cells to maintain epidermal integrity (Donati and Watt 2015). Upon skin wounding, KCs, mainly supplied from hair follicle stem cells, migrate to fill the skin gap (Ito et al. 2005; Page et al. 2013; Donati et al. 2017; Fujimura et al. 2021; Brownell et al. 2011; Joost et al. 2018; Levra Levron et al. 2023). They do so by transforming themselves into EMT phenotypes and stratifying to re-epithelialize through MET when KCs cover the whole wound area (Dongre and Weinberg 2019; Stone et al. 2016; Cheng et al. 2016).

Cell migration into interstitial tissues, namely the extracellular matrix (ECM), is a key feature of wound healing and cancer invasion (Wolf et al. 2013). In this setting, the cells must overcome physical constraints and deform themselves to pass through small gaps, possibly through EMT. These characteristics of cell migration upon spatial confinement have been largely explored using cancer cells in the context of cancer invasion (Wolf et al. 2013; Zulueta-Coarasa et al. 2022; Denais et al. 2016; Mistriotis et al. 2019; Tong et al. 2012). However, if and how non-cancerous epithelial cells migrate into confined spaces is unclear.

Cultured epidermal KCs stratify upon air exposure. This three-dimensional (3D)-reconstructed epidermis (hereafter referred to as 3D epidermis) has been used for disease models and drug screening (Enjalbert et al. 2020; Niehues et al. 2018). The 3D epidermis is typically maintained on a microporous membrane with a pore size of 0.4 µm, which allows medium supply while exposing the cells to air (Sun et al. 2015; Kumamoto et al. 2018, 2021). This model primarily represents steady-state epidermis, although 3D epidermis models have also been applied to wound healing and re-epithelialization studies (Nanes et al. 2024; Kandyba et al. 2008; Sorensen 2006). Wound-related cell migration has additionally been investigated using complementary experimental systems, including the Boyden chamber assay, the scratch-wounding assay (Grada et al. 2017; Guy et al. 2017), and microfluidic migration assays (Sala et al. 2022). However, because these assays do not reflect the 3D nature of epithelial tissues, especially stratified epithelia such as the epidermis, a better experimental system that reproduces skin wounds and their repair is warranted.

We introduce here a novel three-compartment epithelial structure by employing KCs cultured on a 3.0-µm-pored membrane. The tissue model comprises three epithelial stages: a primary stratified epithelium, KCs undergoing EMT, and a newly formed stratified epithelium through MET. Our model recaptures KC migration in physiological wound repair. Through this model, we show that KCs move towards such micropores in an oscillatory manner, which is suppressed by Piezo1 and keratin 6.

## Results

### 3.0-µm micropores induced the formation of a three-compartment epithelial structure

We began with 3D epidermis and questioned whether the size of the membrane pores affects the integrity of the epidermis. To answer this, we cultured primary human epidermal KCs on membranes with different pore sizes smaller than the diameter of the cells (typically 10–20 µm; (Barrandon and Green 1985) and generated 3D epidermis (**Fig. 1a, b**). KCs passed through pores with a diameter of 3.0 µm, which is much smaller than the cells (**Fig. 1b, c**), in agreement with a previous report using other cells (Wolf et al. 2013). Intriguingly, while stratified KCs were maintained above the 3.0-µm-pored membrane, a new stratified epithelium formed under the membrane (**Fig. 1b, c**). Membranes with pores larger than 3.0 µm also allowed KCs to pass through, but the stratified epithelium above the membrane was not well-formed (**Fig. 1b**), contrasting with KCs on the 3.0-µm-pored membrane (**Fig. 1b, c**). KCs within and adjacent to 3.0-µm or larger pores expressed vimentin (Vim), a mesenchymal marker, suggesting that the cells undergo EMT (**Fig. 1b, d, e**). We focused on the 3D epidermis kept on the 3.0-µm-pored membrane because of its distinct phenotype.

**Figure 1.**
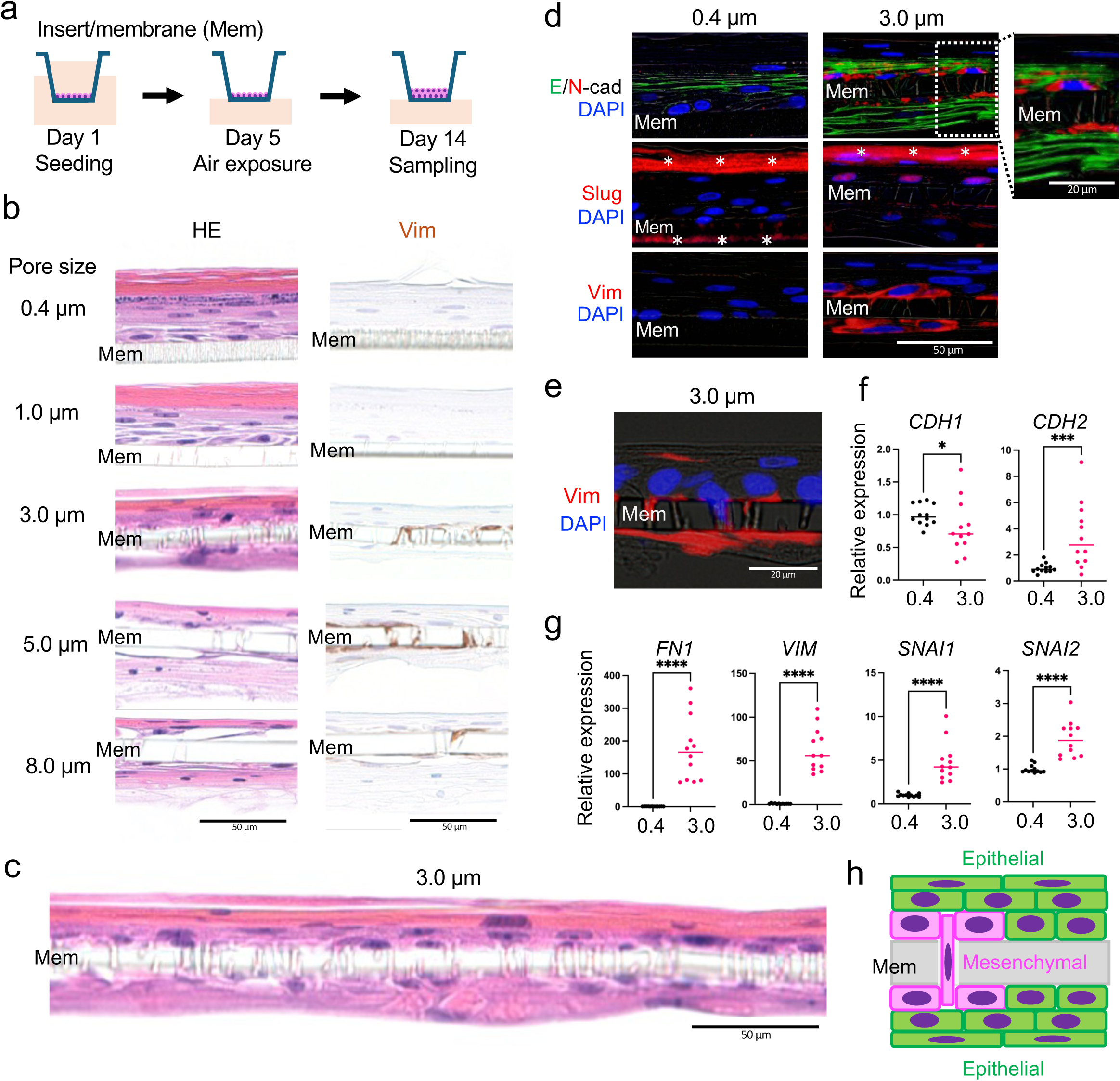
A three-compartment epithelial structure formed by a 3.0-µm-pored membrane. **(a)** Schematic diagram of experiments using normal human epidermal keratinocytes cultured on a microporous membrane and exposed to air for 14 days to produce 3D epidermis. **(b)** Hematoxylin and eosin (HE) staining and vimentin (Vim: brown) labeling of 3D tissue derived from each microporous membrane (Mem). Scale bar: 50 μm. **(c)** HE staining of 3D tissue formed by a 3.0-µm-pored membrane. Scale bar: 50 μm. **(d)** E-cadherin (E-cad)/N-cadherin (N-cad) (upper images), Slug (middle images), and Vim (lower images) staining of 3D tissue formed by a 0.4-µm-(left) or 3.0-µm-pored (right) membrane. Scale bar: 50 μm (low-magnified images) and 20 μm (high-magnified image). **(e)** Vim staining of 3D tissue formed by a 3.0-µm-pored membrane. Scale bar: 20 μm. **(f)** Quantitative RT-PCR (qRT-PCR) of *CDH1* and *CDH2*. **(g)** qRT-PCR of *FN1*, *VIM*, *SNAI1*, and *SNAI2*. N = 12 biological replicates per group. Data were analyzed with two-tailed Mann–Whitney U tests. * p < 0.05; *** p < 0.001; **** p < 0.0001. **(h)** Schematic diagram of the three-compartment epithelial structure. Representative images from three or more replicates per group are shown.

The stratified epithelium above the membrane expressed E-cadherin (E-cad), an epithelial marker, whereas E-cad was reduced in the cells adjacent to the pores, where N-cadherin (N-cad) expression was present (**Fig. 1d**). Instead, Vim and Slug-positive cells, suggestive of EMT, were observed near the pores (**Fig. 1d, e**). The stratified epithelium under the membrane was positive for E-cad again (**Fig. 1d**), implying that the cells retained an epithelial phenotype (MET). Protrusion of KCs within 3.0-µm micropores was also observed (**Fig. 1e**). Quantitative analysis of the immunofluorescence data showed that E-cad expression did not differ significantly among the epithelium on the 0.4-µm-pored membrane and the compartments above and below the 3.0-µm-pored membrane (**Figure 1—figure supplement 1a**). Vim was significantly more abundant in the epithelial structures above and below the 3.0-µm-pored membrane than in those on the 0.4-µm-pored membrane (**Figure 1— figure supplement 1c**). N-cad and Slug showed a similar trend, although the differences did not reach statistical significance except for N-cad (0.4 vs. below 3.0). N-cad was also more abundant below than above the 3.0-µm-pored membrane (**Figure 1—figure supplement 1b, d**). Consistent with the morphological data, gene expression of mesenchymal markers, including *FN1*, *VIM*, *SNAI1*, *SNAI2*, and *CDH2*, was upregulated in the tissue maintained on the 3.0 µm-pored membrane compared with that kept on the 0.4-µm-pored membrane, whereas that of *CDH1*, encoding E-cad, was downregulated in the 3.0-µm-pored group (**Fig. 1f, g**). Laminin β3, a subunit of laminin 332 and a major extracellular matrix component produced by keratinocytes, was detected in the epithelium formed on the 0.4-µm-pored membrane, as well as in all three compartments of the tissue formed on the 3.0-µm-pored membrane (**Figure 1—figure supplement 1e**). These data indicate that the 3.0-µm-pored membrane allowed KCs to form a three-compartment epithelial structure comprising a primary stratified epithelium above the membrane, KCs undergoing EMT within micropores, and another stratified epithelium under the membrane following MET (**Fig. 1h**).

### KCs exhibited oscillatory movement through 3.0-µm micropores

We then asked how the three-compartment epithelial structure developed. To answer this, we established HaCaT KCs stably expressing GFP-LifeAct, which labels F-actin in living cells (hereafter referred to as LifeAct-KCs; **Figure 2—figure supplement 1a**). Aligning with the results of primary KCs (**Fig. 1**), LifeAct-KCs protruded into 3.0-µm micropores with filopodia-like structures, but not into 0.4- or 1.0-µm micropores (**Figure 2—figure supplement 1b**). Electron microscopy also showed that LifeAct-KCs extended their cytoplasm into the micropores (**Figure 2— figure supplement 1c**). To further investigate the dynamics of the cells, we conducted live imaging using LifeAct-KCs. Just below the 3.0-µm-pored membrane, cells that had exited the pores and expanded on the bottom of the membrane were observed (**Fig. 2a, b**; arrows, **Supplementary Movie 1**), but at the same time, some cells re-entered the pores (**Fig. 2a, b**; arrowheads, **Supplementary Movie 1**). This oscillatory movement of filopodia-like protrusions was more visible when the focus was set at the level of the micropores, and these protrusions were extended into and out of the pores (**Fig. 2c, d, Supplementary Movie 2**). We wondered whether this oscillatory movement of filopodia-like protrusions was specific to the cells kept on 3.0-µm micropores or universal in migrating cells on microporous membranes. Whereas LifeAct-KCs kept on the 3.0-µm-pored membrane showed oscillatory movement of filopodia-like protrusions, the cells kept on 8.0-µm micropores were prone to unidirectional movement (**Fig. 2e, f, Supplementary Movie 3, 4**), which was confirmed by quantitative analysis (**Fig. 2g, h**, see Materials and Methods section). In line with the data on the three-compartment epithelial structure (**Fig. 1c**), LifeAct-KCs above the membrane and within the micropores expressed both E-cad and Vim, showing at least a partial EMT phenotype (**Fig. 2i, j**). The Vim/E-cad ratio was significantly lower within the micropores than above the membrane (**Figure 2— figure supplement 1d**), indicating that the partial EMT phenotype was already present in the cells above the membrane rather than being acquired upon entry into the micropores. These data indicate that the KCs kept on 3.0-µm micropores initially move in an oscillatory manner (**Fig. 2e, g, h, Supplementary Movie 3**) but can move out of the pores to form a stratified epithelium under the membrane (**Fig. 2b**), possibly accounting for the formation of the three-compartment epithelial structure (**Fig. 1b, c**). In contrast, the unidirectional movement of KCs through 8.0-µm micropores (**Fig. 2f–h, Supplementary Movie 4**) might explain the improperly formed epithelium above the 8.0 µm-pored membrane (**Fig. 1b**).

**Figure 2.**
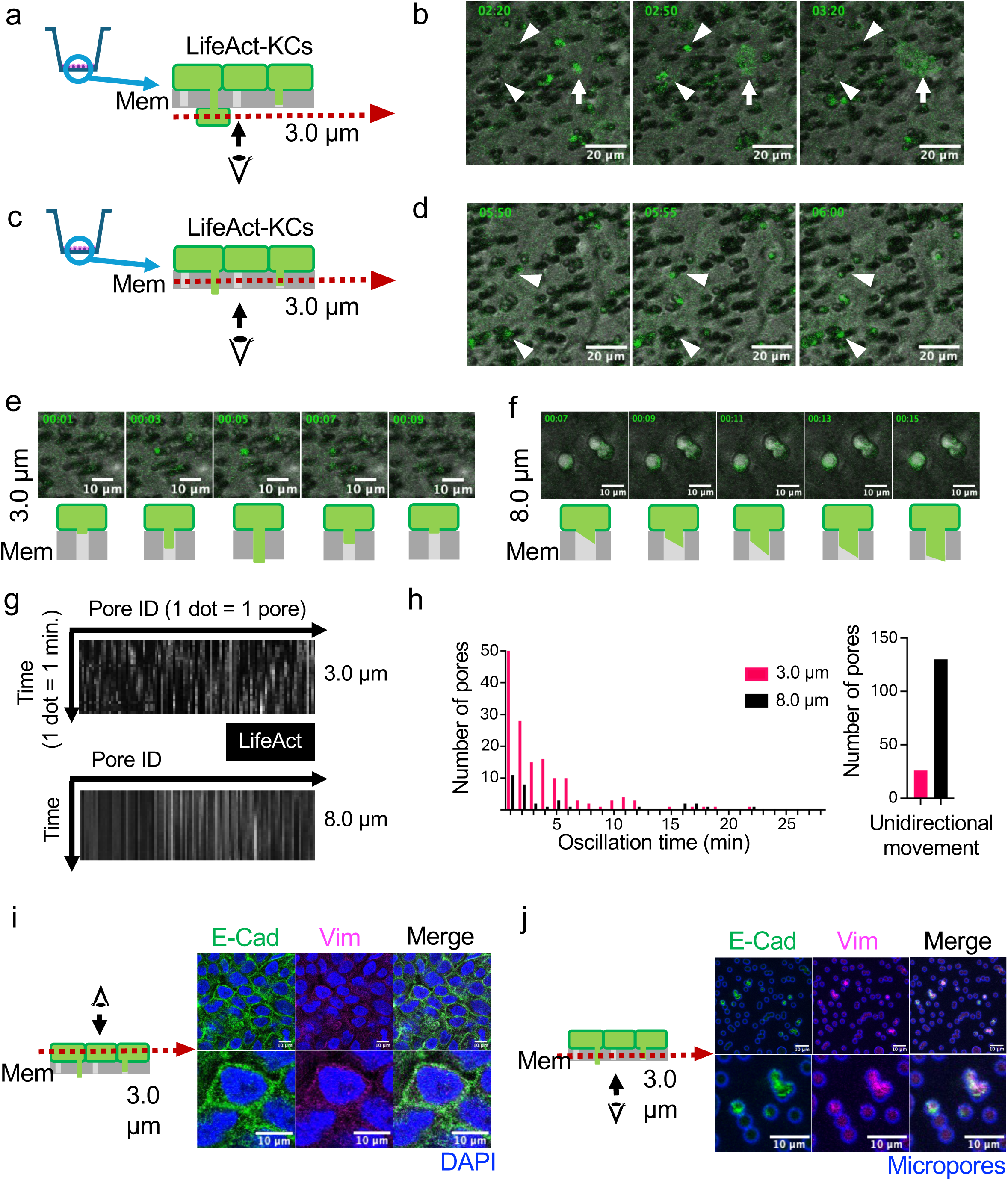
Oscillatory movement of keratinocytes on a 3.0-µm-pored membrane. **(a)** Schematic diagram of LifeAct-KCs cultured on a microporous membrane. Observation at the level of cell exit from the micropores. **(b)** Time-lapse images of LifeAct-KCs at the level of cell exit from 3.0-µm micropores. Representative images are shown at 30-minute time intervals. Time stamp shows hour:minute. Arrows indicate the cells that exit from the pores and expand under the membrane. Arrowheads indicate the cells that show oscillatory movement. Scale bar: 20 µm. **(c)** Schematic diagram of LifeAct-KCs cultured on a microporous membrane. Observation at the level of cell entry into the micropores. **(d)** Time-lapse images of LifeAct-KCs at the level of cell entry into 3.0-µm micropores. Representative images are shown at 5-minute intervals. Time stamp shows hour:minute. Arrowheads indicate the cells that show oscillatory movement. Scale bar: 20 µm. **(e, f)** Time-lapse images and schematic diagrams of LifeAct-KCs at the level of cell entry into 3.0-µm **(e)** or 8.0-µm **(f)** micropores. Representative images are shown at 2-minute intervals. Time stamp shows hour:minute. Scale bar: 10 µm. **(g, h)** Quantification of oscillatory movement of LifeAct-KCs on 3.0-µm or 8.0-µm micropores. **(g)** LifeAct signals (white) are shown as a spectrogram, where the X-axis represents each pore, and the Y-axis represents time. Data are summarized from three independent experiments. **(h)** Graphs comparing the number of pores presenting the LifeAct signals within 3.0-µm or 8.0-µm micropores per time. The left panel shows oscillation time as the X-axis and the number of pores as the Y-axis. The right panel shows unidirectional movement of the cells (no oscillation within 30 minutes) within each micropore. Data are summarized from three independent experiments. **(i, j)** E-cad and Vim labeling of LifeAct-KCs at the level of cells above **(i)** or within **(j)** 3.0-µm micropores. Autofluorescence of micropores is shown in blue (**j**). Scale bar: 10 µm. Representative images from three or more replicates per group are shown.

### Actin polymerization is essential for KC migration through 3.0-µm micropores

We next investigated the mechanisms underlying KC migration through 3.0-µm micropores, which involves two major steps: cell entry into the gaps and cell exit from the pores. We first examined the TGF-β pathway because KCs on the micropores showed EMT phenotypes (**Fig. 1b–d**), and the activation of the TGF-β pathway is associated with EMT (Kalluri and Weinberg 2009). TGF-β ligand promoted LifeAct-KC entry into 3.0-µm micropores (**Fig. 3a**, **Figure 3—figure supplement 1a**), but blocking TGF-β receptors did not affect this process (**Fig. 3b**, **Figure 3—figure supplement 1b**). The cell exit from the micropores showed similar results across treatments (**Fig. 3d, e**, **Figure 3—figure supplement 1d, e**). We then looked into actin because it was enriched in the actin-rich filopodia-like protrusions extending into 3.0-µm micropores (**Fig. 2b**). As expected, the inhibition of actin polymerization with cytochalasin D prevented KC entry into 3.0-µm micropores (**Fig. 3c**, **Figure 3—figure supplement 1c**) and slowed cell exit from the micropores (**Fig. 3f**, **Figure 3—figure supplement 1f**). Overall, these data suggest that KC migration through confined spaces is promoted by, but does not require, TGF-β and that this migration depends on actin polymerization.

**Figure 3.**
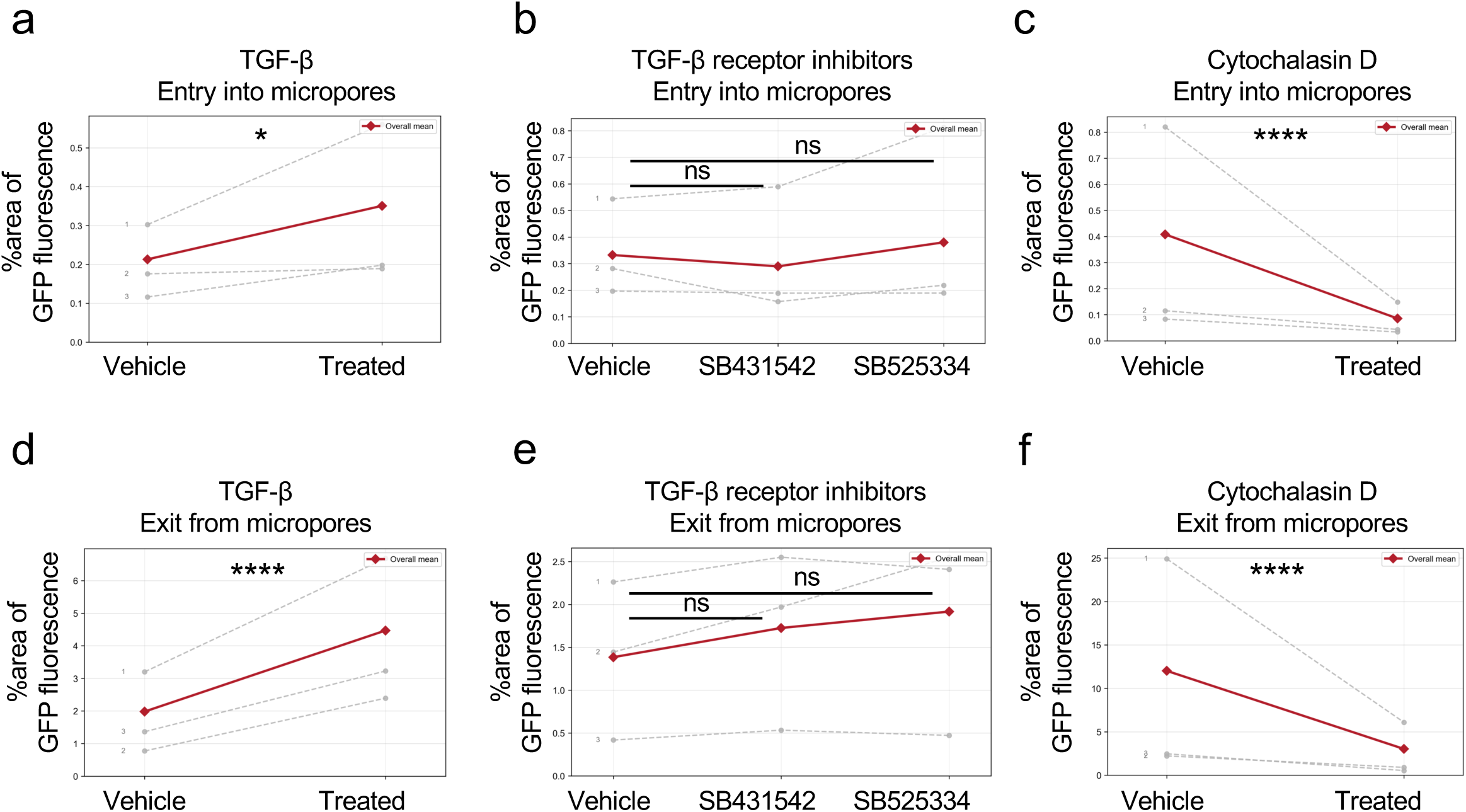
Chemical treatment on KC entry into and exit from 3.0-µm micropores. **(a–c)** Quantification of LifeAct-KCs present within 3.0-µm micropores 6 hours after cell seeding. Cells were treated with TGF-β ligand **(a)**, TGF-β receptor inhibitors, SB431542, and SB525334 **(b)**, or cytochalasin D **(c**). **(d–f)** Quantification of LifeAct-KCs present below 3.0-µm micropores 24 hours after cell seeding. Cells were treated with TGF-β ligand **(d)**, TGF-β receptor inhibitors **(e)**, or cytochalasin D **(f**). Each dotted line represents an experimental batch, and the red line indicates the overall mean. N = 7 biological replicates per group. Data were analyzed using a gamma generalized linear model with a log link function, adjusting for experimental batch as a categorical fixed effect. * p < 0.05; **** p < 0.0001; ns, not significant.

### Piezo1 and keratin 6 hinder KC migration through 3.0-µm micropores and 3D cell invasion

We then hypothesized that mechanosensors might affect KC migration through 3.0-µm micropores. Intriguingly, ruthenium red, a universal ion-channel blocker, promoted LifeAct-KC entry into 3.0-µm micropores and showed a similar, albeit statistically non-significant, trend for exit (**Fig. 4a, b**, **Figure 4—figure supplement 1a, b**). Piezo1 is a major mechanosensitive ion channel in KCs, the inhibition of which was implicated in speeding KC migration in a scratch-wounding assay (Holt et al. 2021). Piezo1 knockout (PIEZO1 KO) HaCaT KCs showed accelerated LifeAct-KC entry into and exit from 3.0-µm micropores (**Fig. 4c, d**, **Figure 4—figure supplement 1e, f, i**), whereas Piezo1 agonist Yoda1 slowed these processes (**Fig. 4e, f**, **Figure 4—figure supplement 1c, d**).

**Figure 4.**
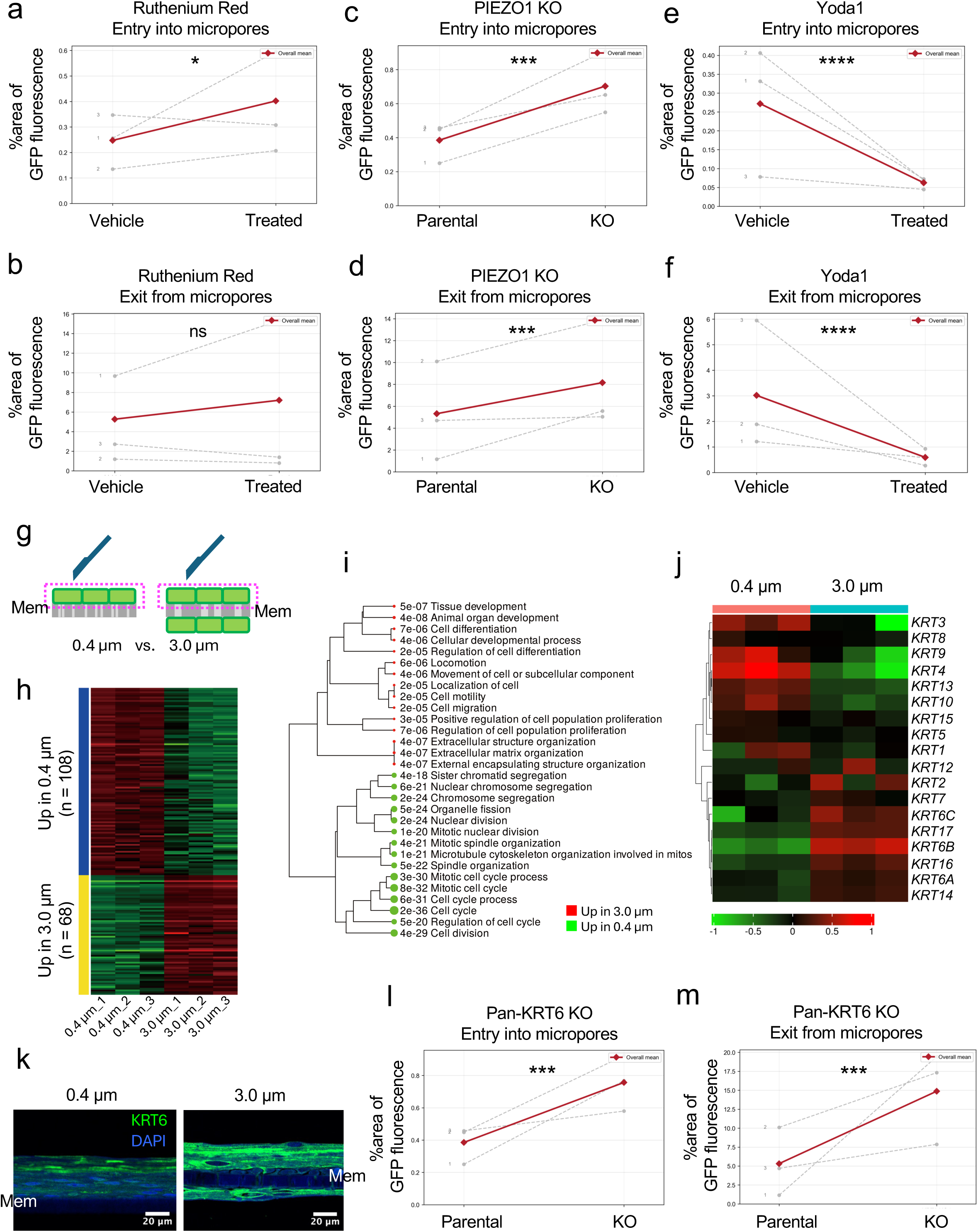
Regulatory role of Piezo1 and KRT6 on KC entry into and exit from 3.0-µm micropores. **(a, b)** Quantification of ruthenium red–treated LifeAct-KCs present within 3.0 µm micropores at 6 hours **(a)** and the cells present below 3.0-µm micropores 24 hours after cell seeding **(b)**. **(c, d)** Quantification of PIEZO1 KO LifeAct-KCs present within 3.0 µm micropores at 6 hours **(c)** and the cells present below 3.0-µm micropores 24 hours after cell seeding **(d)**. **(e, f)** Quantification of Yoda1-treated LifeAct-KCs present within 3.0-µm micropores at 6 hours **(e)** and the cells present below 3.0-µm micropores 24 hours after cell seeding **(f)**. Each dotted line represents an experimental batch, and the red line indicates the overall mean. N = 7 **(a, b)** or 9 **(c–f)** biological replicates per group. Data were analyzed using a gamma generalized linear model with a log link function, adjusting for experimental batch as a categorical fixed effect. * p < 0.05; *** p < 0.001; **** p < 0.0001; ns, not significant. **(g)** Schematic diagram of bulk RNA-seq comparing HaCaT KCs cultured on a 0.4-µm- or 3.0-µm-pored membrane. The magenta dashed rectangles indicate the cells compared in the bulk RNA-seq. **(h)** Heatmap of differentially expressed genes identified by the bulk RNA-seq. The full gene lists are provided in Figure 4—source data 1. **(i)** Gene ontology biological process terms enriched in upregulated and downregulated genes. **(j)** Heatmap of differentially expressed keratin genes in HaCaT KCs cultured on a 0.4-µm- or 3.0 µm-pored membrane. **(k)** Keratin 6 (KRT6) labeling of normal human epidermal KCs cultured on a 0.4-µm- or 3.0-µm-pored membrane for 14 days. Scale bar: 20 µm. Representative images from three replicates per group are shown. **(l, m)** Quantification of pan-KRT6 KO LifeAct-KCs present within 3.0-µm micropores at 6 hours **(l)** and the cells present below 3.0-µm micropores 24 hours after cell seeding **(m)**. Each dotted line represents an experimental batch. N = 9 biological replicates per group. The parental data are shared between panels c, d and l, m, because the parental, PIEZO1 KO, and pan-KRT6 KO cell lines were assayed in parallel in the same experiments. Data were analyzed using a gamma generalized linear model with a log link function, adjusting for experimental batch as a categorical fixed effect. *** p < 0.001.

To further delineate the mechanisms underlying this phenomenon, we performed bulk RNA-seq for LifeAct-KCs above 0.4- and 3.0-µm micropores (**Fig. 4g–i**, **Figure 4—source data 1**). The gene ontology biological process terms enriched among the differentially expressed genes (DEGs) through DESeq2 (Love et al. 2014) included tissue development and cell differentiation (**Fig. 4i**). Keratins are major cytoskeletal proteins, and differential keratin expression accounts for KC development and differentiation. Among keratin genes, *KRT6A*, *KRT6B*, and *KRT6C*, all encoding keratin 6 (a stress-induced keratin), were upregulated in the KCs kept on 3.0-µm micropores (**Fig. 4j**). Keratin 6 was also pronounced in the three-compartment epithelial structure derived from normal human epidermal KCs (**Fig. 4k**). To uncover the role of *KRT6A*, *KRT6B*, and *KRT6C*, we generated HaCaT KCs with KO for these three keratins (pan-KRT6 KO KCs; **Figure 4—figure supplement 1g, h**). Pan-KRT6 KO KCs showed pronounced cell entry into and exit from 3.0-µm micropores (**Fig. 4l, m**, **Figure 4—figure supplement 1i**). These data indicate that Piezo1 and keratin 6 negatively modulate KC migration through confined spaces.

## Discussion

Our study presents a three-compartment epithelial structure in which the original epithelium above 3.0-µm micropores and a new epithelium under the micropores were interconnected with non-cancerous KCs exhibiting EMT. The 3.0-µm micropores specifically induced the unique epithelial tissue because larger micropores failed to maintain the epithelium above the membrane and smaller pores did not allow the cells to pass through. KCs entered the 3.0-µm micropores but showed oscillatory movement until they finally exited the pores. This oscillatory movement (micropore-induced oscillatory movement; MIOM) might underlie the formation of the three-compartment epithelial structure because, otherwise, the unidirectional migration of KCs toward the exit from the pores could deplete the epithelium above the micropores.

Although the TGF-β ligand accelerated the KC entry into 3.0-µm micropores, the TGF-β pathway is dispensable because the blockade of TGF-β receptors did not slow this process. This is probably because the pivotal step of KC entry into 3.0-µm micropores involves the extension of actin-rich filopodia-like structures into the micropores, which might not be regulated by the TGF-β pathway. As expected, inhibiting actin polymerization effectively prevented KC entry into 3.0-µm micropores. Reduced Piezo1 is known to accelerate wound healing in mice and KC migration in 2D cultures (Holt et al. 2021). This is explained by the role of Piezo1 in regulating cell shape, causing cell retraction (Holt et al. 2021) and suppressing leader cell formation (Chen et al. 2024). Our data concerning the negative impact of Piezo1 on cell entry into 3.0-µm micropores might be associated with the less polarized shape of Piezo1-null KCs, as described previously (Holt et al. 2021). Since ruthenium red is a universal ion-channel blocker, mechanosensors other than Piezo1 may be involved in cell entry into micropores. Further research is required to address this issue.

Keratin 6 is a stress-induced keratin that interacts with keratin 16 and 17 (Cohen et al. 2024). Monoallelic variants in *KRT6A*, *KRT6B*, and *KRT6C*, as well as *KRT16* and *KRT17*, are causal for pachyonychia congenita, a disease in which patients develop nail dystrophy, steatocystoma multiplex, and calluses (O’Toole et al. 2024). *Krt6a-* and *Krt6b*-null KCs have shown faster collective cell migration by regulating myosin ATPase activity (Wang et al. 2018), which is possibly compatible with the accelerated cell exit from 3.0-µm micropores of pan-KRT6 KO KCs in our study. In addition, as exemplified by the increase of stress fibers in *Krt6a*- and *Krt6b*-null KCs (Wong and Coulombe 2003), more direct effects on actin organization in pan-KRT6 KO KCs may be involved in the cell entry into the micropores.

The mechanistic relationship among actin polymerization, keratin 6, and Piezo1 in regulating KC migration through confined spaces remains to be fully elucidated. Our data support a model in which mechanosensing and cytoskeletal remodeling, rather than transcriptional EMT programs alone, primarily govern KC migration through micropores. Actin polymerization appears to be the central driver of cell entry into and exit from the pores, whereas Piezo1 and keratin 6 likely influence this process by modulating cytoskeletal organization and cellular mechanical properties.

HaCaT keratinocytes were used in this study primarily because of their suitability for genetic manipulation and stable expression of fluorescent reporters. We acknowledge that HaCaT cells, as an immortalized keratinocyte line, do not recapitulate all features of primary human epidermal keratinocytes, including aspects of differentiation, genomic stability, and cell migration (Pilcher et al. 1997). However, key observations, including migration into 3.0-µm micropores and the formation of the three-compartment epithelial structure, were initially made in primary keratinocytes. The use of HaCaT cells therefore enabled mechanistic interrogation while maintaining consistency with findings in primary cells.

Wound healing requires several components and steps: an intact surrounding epithelium that supplies migrating cells to heal the wound, KC migration to the wound area, and re-epithelialization once the wound is fully covered. The three-compartment epithelial structure in our study reproduces these components in a culture system, and the advantage of this model is that each step of the wound healing process is simultaneously visible. Another idea is that the micropores in our system may simulate holes in the ECM or basement membrane (BM), which epithelial cells protrude into during migration. The intact BM pores are estimated to be around 10 nm (Yurchenco 2011), much smaller than the micropores in our study. However, in special circumstances such as wound healing, BM can be microbreached due to physical insults or proteases released from KCs or other cells. Similar cell migration occurs in salivary gland development, in which epithelial cells protrude into BM holes with a size of µm (Harunaga et al. 2014). Aligning with our study, salivary epithelial cells move in and out of BM during development (Harunaga et al. 2014).

Although Piezo1, TGF-β signaling, and cytoskeletal dynamics have previously been implicated in KC migration, the conceptual advance of our study lies in demonstrating how confined spatial architecture alone can generate a spatially organized epithelium comprising spatially graded EMT and MET states within a single continuous tissue. In our system, KCs undergoing migration through confined spaces exhibited oscillatory protrusive behavior that preserves the integrity of the overlying epithelium while simultaneously enabling the formation of a new stratified epithelium beneath the membrane. This dynamic integration of migration and EMT/MET within a stratified epithelial context has not, to our knowledge, been previously demonstrated. The oscillatory migratory mode observed here differs from the predominantly unidirectional migration described in conventional two-dimensional or Transwell assays and suggests that physical confinement can actively shape epithelial organization and migratory dynamics in a self-organizing manner.

In conclusion, our study presents a three-compartment epithelial structure in a culture system, which is possibly based on the unique phenomenon of MIOM. Our model can be exploited to reproduce wound healing and epithelial development and help develop therapeutics for skin wounds.

## Methods

### Cells

Normal human epidermal KCs collected from neonatal foreskin were purchased from Kurabo. HaCaT KCs, originally derived from the back skin of a 62-yr-old man (Boukamp et al. 1988), were provided by Dr. Norbert Fussenig’s lab (German Cancer Research Center), and the cell identity was confirmed with the Cell Culture STR profile (Biologica). Single-cell-cloned HaCaT KCs with stable Cas9 expression were established previously to enable later KO experiments (Mai et al. 2023, 2024). The GFP-LifeAct-expressing vector was generated by inserting the LifeAct sequence (a gift from Dr. Kurisu, Tokushima University) into the pRetroQ-AcGFP1-C1 vector (Takara Bio). The GFP-LifeAct-expressing vector was transfected into Ampho-Pack 293 cells (Clontech) using Avalanche Everyday Transfection Reagent (EZ Biosystems) according to the manufacturer’s instructions. GFP-LifeAct-expressing retrovirus was collected from the culture medium of transfected Ampho-Pack 293 and infected into Cas9-expressing HaCaT KC with 8 µg/mL polybrene (Sigma-Aldrich). After the infection, GFP-LifeAct-expressing cells were sorted by BD FACSAria III (BD Biosciences) and single-cell cloned to generate LifeAct-KCs.

PIEZO1 KO KCs and pan-KRT6 KO KCs were established from LifeAct-KCs as described previously (Mai et al. 2024). CRISPR RNA (crRNA) was acquired from predesigned Alt-R CRISPR-Cas9 guide RNA (Integrated DNA Technologies), and the crRNA contained the CACGCGCTGGTCCTCAACACCGG sequence, with the CGG PAM sequence targeting exon 8 of the *PIEZO1* genomic sequence (NG_042229.1). The crRNA to generate pan-KRT6 KO KCs contained the CGACCTGCGCAACACCAAGCAGG sequence to target a shared region among *KRT6A* (NG_008298.1), *KRT6B* (NG_008299.1), and *KRT6C* (NG_012416.1). The KO of each gene was confirmed by quantitative RT-PCR (qRT-PCR) and Western blotting. We routinely tested all cells for mycoplasma contamination and confirmed negative results.

### Cell culture

Normal human epidermal KCs were thawed and cultured in EpiLife Medium with 60 µM calcium (ThermoFisher) using a supplement kit containing insulin, human epidermal growth factor, hydrocortisone, antibiotics, and bovine pituitary extract (Kurabo), then stocked in CELLBANKER 2 (ZNQ) after two passages. For NHEK 3D modeling, NHEKs were thawed from stocks into EpiLife, treated with trypsin at subconfluent, and suspended in CnT-Prime (CELLnTEC) at a concentration of 2.5x10^5^ cells/mL for seeding. The day before seeding, Millicell Hanging Cell Culture Insert, PET 0.4, 1.0, 3.0, 5.0, or 8.0 µm, 12-well (Merck), was placed in 12-well plates (Corning) and coated with CELLstart solution (ThermoFisher) diluted 50-fold in PBS with MgCl_2_ and CaCl_2_ (Sigma-Aldrich) and incubated overnight at 4 °C. After removal of the coating solution, 1 mL of CnT-Prime was added in 12 well plates, and 1 mL of cell suspension was seeded inside the inserts. Three days after seeding, the medium was replaced with CnT-Prime 3D Barrier Culture Medium (CELLnTEC) supplemented with L(+)-ascorbic acid at a concentration of 50 µg/mL. The medium inside the insert was removed, and the cell surface was exposed to air on the following day. After the air exposure, Extra Thick Blot Filter Paper (Bio-Rad), cut into 28 mm squares and with two 12-mm holes, was placed on Falcon 6-well deep well plates (Corning), and each well was filled with 11.5 mL of CnT-Prime-3D (Mai et al. 2024). The inserts were placed on the holes of the filter paper, followed by medium changes every 3 days. After 14 days from seeding, the membrane was cut from the insert and used for histology, immunofluorescence, and RNA extraction.

Cas9-expressing HaCaT KCs and LifeAct-KCs were cultured in Dulbecco’s modified eagle medium (DMEM) with 4.5g/L glucose and L-glutamine (Nacalai Tesque, Kyoto, Japan) supplemented with 10% fetal bovine serum (FBS; HyClone) and 1x antibiotic–antimycotic mixed stock solution (Nacalai Tesque). The procedure for 3D modeling of HaCaT KCs using inserts was similar to that used for normal human epidermal KCs, except that the DMEM, FBS, and antibiotics were used after seeding and air exposure.

### Live cell imaging

Live cell imaging on LifeAct-KCs cultured on a microporous membrane was carried out with LSM710 (Carl Zeiss) or FV1000 (Olympus) confocal microscopes equipped with an incubation chamber maintained at 37 °C and 5% CO_2_. During live cell imaging, LifeAct-KCs were cultured in DMEM (4.5 g/L glucose) without phenol red (Nacalai Tesque) supplemented with L-Glutamine (ThermoFisher). LifeAct-KCs were seeded at 7.0x10^5^ cells/mL per pre-coated insert. The inserts were placed on the glass-bottomed dishes containing 1 mL of medium. The lids were fixed by wrapping their sides with Parafilm with air holes. Live cell imaging was performed after 6 hours of incubation from seeding.

### Quantification of oscillatory movement of KCs

The time sequence of the image files taken with the LSM710 and FV1000 was used for the quantification of the cell movement. Cellular locations were estimated by the region with high GFP intensity. We first smoothed the images by taking the average intensity of the nine neighboring pixels. Then, the pixels with top *X* percent intensity were detected. If more than *Y* pixels with top *X* percent high-intensity pixels were connected, the region should correspond to the pore where a cell entered and exited. The pore region was defined as a circular region with a radius of *Z* pixels, where the center corresponds to the center of mass of the connected region. We calculated the average intensity in each pore region as a function of time. For each pore region, the minimum of the maximum values in each pore region of one experiment was used as the threshold, and the number of consecutive images exceeding the threshold was counted. When the first or last image exceeded the threshold, the consecutive counts were excluded if it was 5 or less. We set the parameters *X* = 1, *Y* = 10, and *Z* = 5 for the case with 3-μm pores and *X* = 3, *Y* = 50, and *Z* = 20 for the case with 8-μm pores. Three independent experiments were performed, and the pooled data are presented.

### Quantification of cell entry into and cell exit from micropores

Image files taken with the LSM710 and FV1000 were uniformly processed by an ImageJ macro that binarized the images after Gaussian blurring. For each image, the %Area was calculated by dividing the number of GFP pixels by the total number of pixels in each image.

### Ligand and chemical treatments

Recombinant human TGF-β1 (Peprotech), TGF-β receptor 1 inhibitor, SB431542 (Selleck) and SB525334 (Selleck), cytochalasin D (Cayman Chemical), ruthenium red (Wako), and Yoda1 (Cayman Chemical) were used at concentrations of 5 ng/mL, 10 µM, 10 µM, 0.25 µM, 100 µM, and 25 µM, respectively. These reagents were added at seeding, except cytochalasin D and Yoda1, which were added 5.5 hours after seeding because they prevent cell attachment to the membrane. The cells were evaluated 6 hours after seeding for cell entry into the micropores and at 24 hours for cell exit from the pores after seeding. Cytochalasin D was washed out with fresh medium after the imaging, 6 hours after seeding.

### RNA extraction and quantitative RT-PCR

RNA was extracted from normal human epidermal KCs or cultured HaCaT cells using the RNeasy Mini kit (Qiagen, Hilden, Germany). cDNA was prepared using the SuperScript IV First-Strand Synthesis System (Thermo Fisher Scientific). qRT-PCR was performed using the designated primers and fast SYBR Green (Thermo Fisher Scientific) in a STEP-One plus sequence detection system (Applied Biosystems, Waltham, MA, USA). All primers used in this study are listed in Supplementary Table 1. Pan-*KRT6* primers were designed based on the sequences common to *KRT6A*, *KRT6B*, and *KRT6C*. *GAPDH* and *YWHAZ* were used as internal controls in the qRT-PCR of normal human epidermal KCs and HaCaT KCs, respectively.

### Western blotting

Cell lysates were collected using a lysis buffer containing 1% Nonidet P-40 (Nacalai Tesque), 25 mM Tris–HCl (pH 7.6), 100 mM NaCl, 10 mM ethylenediaminetetraacetic acid, and a 1:100 dilution protease inhibitor cocktail (P8340; Sigma-Aldrich) on ice for 30 minutes with shaking. The lysates were centrifuged at 15,300 g at 4 °C for 20 minutes. Cell lysate supernatants were denatured with a 5× loading buffer (0.25 M Tris–HCl; 8% sodium dodecyl sulfate; 30% glycerol; 0.02% bromophenol blue; 0.3 M β-mercaptoethanol; pH 6.8). Samples were subjected to electrophoresis using NuPAGE 4% to 12%, Bis-Tris, 1.0–1.5 mm, Mini Protein Gels (Thermo Fisher Scientific). Proteins separated in the gels were transferred onto PVDF transfer membranes (Bio-Rad). The membranes were blocked with 2% skim milk and incubated with rabbit anti-PIEZO1 antibody (Cat# NBP1-78537, RRID:AB_11003149, 1:500 dilution; Novus), rabbit anti-KRT6A antibody (Cat# PRB-169P, RRID:AB_10063923, 1:1,000 dilution; Biolegend), or mouse anti-GAPDH antibody (Cat# G8795, RRID:AB_1078991, 1:5,000 dilution; Sigma) in 2% skim milk at 4 °C overnight. After washing with Tris-buffered saline, the membranes were incubated with peroxidase-conjugated anti-rabbit IgG antibody (Cat# 711-035-152, RRID:AB_10015282, 1:5,000 dilution; Jackson

ImmunoResearch) or anti-mouse IgM antibody (Cat# 315-035-020, RRID:AB_2340064, 1:5,000 dilution; Jackson ImmunoResearch) in 2% skim milk at RT for 1 hour. Signals were visualized by Clarity Western ECL Substrate (Bio-Rad) and detected with an ImageQuant LAS 4000 mini camera system (Fujifilm).

### Electron microscopy

Ultrastructural observation of HaCaT KCs was performed as previously described (Kosumi et al. 2022). Briefly, HaCaT KCs were cultured for 4 hours after seeding in DMEM supplemented with 10% FBS (HyClone) and 1× antibiotic–antimycotic mixed stock solution (Nacalai Tesque) on 3.0-µm-pored cell culture inserts (Merck KGaA, Darmstadt, Germany). The inserts were pre-coated with CELLstart (Thermo Fisher Scientific, Waltham, MA, USA). HaCaT KCs on the inserts were fixed with 2% glutaraldehyde and 2% paraformaldehyde in 30 mM 4-(2-hydroxyethyl)-1-piperazineethanesulfonic acid buffer. The samples were dehydrated and embedded in Epon 812. Thin sections for electron microscopy were cut to 70 nm in thickness, stained with uranyl acetate and lead citrate, and examined using a JEM1400 (Japan Electron Optics Laboratory, Tokyo, Japan).

### RNA-seq

HaCaT KCs were cultured on a 0.4-µm- or 3.0-µm-pored micromembrane for 48 hours. KCs above the membrane were then collected using a scraper. RNA was extracted using the RNeasy Mini kit (Qiagen, Hilden, Germany). Library preparation, sequencing, and further analysis were performed as described previously (Mai et al. 2024).

### Histology

KCs cultured on a microporous membrane were fixed with formalin and embedded in paraffin after dehydration. Sectioned paraffin samples were deparaffinized and stained with hematoxylin and eosin (HE). For immunohistochemistry, KCs were fixed with 4% paraformaldehyde in PBS, followed by dehydration and embedding in paraffin. Anti-Vim antibody (Cat# EPR3776, RRID:AB_10562134, 1:500 dilution; Abcam), biotin-conjugated anti-rabbit IgG antibody (Cat# E0432, RRID:AB_2313609, 1:200 dilution; Agilent), VECTASTAIN ABC Standard Kit (Vector Laboratories), and DAB substrate (Roche Diagnostics Inc.) were used for staining. Images were taken with a BZ-9000 microscope (Keyence, Osaka, Japan).

### Immunofluorescence

Cultured KCs were fixed with 4% paraformaldehyde, followed by permeabilization with 0.1% Triton X-100 in PBS for 20 minutes at room temperature. Formalin-fixed paraffin-embedded samples were sectioned and deparaffinized. The cells or sectioned deparaffinized samples were then incubated with primary antibodies overnight at 4. After washing in PBS, the cells were incubated with secondary antibodies conjugated with Alexa488, Alexa568, or Alexa647 for 1 hour at room temperature. The nuclei were stained with 4’,6-diamidino-2-phenylindole. All stained immunofluorescent samples were observed using a confocal laser scanning microscope (FV1000, Olympus). The images were analyzed using ImageJ. The following antibodies were used: anti-Vim (Cat# EPR3776, RRID:AB_10562134, 1:500 dilution; Abcam or Cat# sc-373717, RRID:AB_10917747, 1:200 dilution; Santa Cruz), anti-E-cad (Cat# AF648, RRID:AB_355504, 1:1,000 dilution; R and D system or Cat# 3195, RRID: AB_2291471, 1:200 dilution; Cell Signaling), anti-N-cad (Cat# ab18203, RRID:AB_444317, 1:100 dilution; Abcam), anti-SLUG (Cat# 9585, RRID:AB_2239535, 1/100 dilution; Cell Signaling), anti-laminin β3 (Cat# MA5-24654, RRID:AB_2607185, 1:50 dilution; ThermoFisher Scientific), and anti-KRT6A (Cat# PRB-169P, RRID:AB_10063923, 1:500 dilution; Biolegend).

### Statistics

Statistical analyses were performed with GraphPad Prism 10 (GraphPad Software), except for the generalized linear models, which were fitted in Python 3.13.9 using the statsmodels package (version 0.14.5). P-values were determined with two-tailed Mann–Whitney U tests, one-way analysis of variance (ANOVA) followed by Tukey’s multiple comparison test, or paired t-tests, as indicated in each figure legend. P-values of <0.05 were considered statistically significant (* p < 0.05, ** p < 0.01, *** p < 0.001, **** p < 0.0001). In experiments with KO cells and chemical treatments, for each biological replicate, eight measurement points were quantified, and their mean was used as the representative value for that sample. An experimental batch consisted of 2 to 3 biological replicates per group that were processed and imaged in parallel. To adjust for batch-to-batch variability, the experimental batch was included as a categorical fixed effect in the generalized linear model. Treatment was included as the main explanatory variable. Because the outcome variable was continuous and positive, a gamma generalized linear model with a log link function was used: percentage area ∼ treatment + C(experimental_batch), where C(experimental_batch) denotes that the experimental batch was modeled as a categorical variable. This approach adjusted for batch-specific baseline differences without assuming a linear trend across batches.

## Data availability

RNA-seq data have been deposited in GEO under accession number GSE282553 (https://www.ncbi.nlm.nih.gov/geo/query/acc.cgi?acc=GSE282553).

## Supporting information

Figure 3-igure supplement 1

## Acknowledgments

We thank Ms. Mika Tanabe and the Nikon Imaging Center at Hokkaido University for their technical assistance. We also thank Dr. Mitsuhiro Denda for his invaluable input and support. This work was funded by JST CREST (grant number JPMJCR1926) to MN, the Project for Junior Scientist Promotion of Hokkaido University to HU, and JSPS KAKENHI (grant number 23H02928), JST FOREST Program (grant number JPMJFR245F), the Naito Foundation, and the Nakatomi Foundation to KN. The work was also supported by JSPS KAKENHI (grant number JP22H04926) Grant-in-Aid for Transformative Research Areas―Platforms for Advanced Technologies and Research Resources “Advanced Bioimaging Support.”

## Supplementary Figure Legends

**Figure 1—figure supplement 1. Quantitative analysis of EMT marker expression and laminin β3 distribution**

**(a–d)** Mean fluorescence intensity (A.U.) of E-cad **(a)**, N-cad **(b)**, Vim **(c)**, and Slug **(d)** in the epithelium formed on the 0.4-µm-pored membrane and in the compartments above and below the 3.0-µm-pored membrane. N = 3 biological replicates per group. Data were analyzed with one-way ANOVA followed by Tukey’s multiple comparison test. * p < 0.05; ** p < 0.01; *** p < 0.001; ns, not significant. **(e)** Laminin β3 staining of 3D tissue formed by a 0.4-µm-(left) or 3.0-µm-pored (right) membrane. Scale bar: 50 μm. Representative images from four replicates per group are shown.

**Figure 2—figure supplement 1. Characterization of HaCaT KCs and LifeAct-KCs**

**(a)** LifeAct-KCs kept on a dish. Scale bar: 20 µm. **(b)** HE staining of parental HaCaT KCs cultured on a 1.0- or 3.0-µm-pored membrane for 14 days. Scale bar: 20 µm. **(c)** Ultrastructural findings of HaCaT KCs cultured on a 3.0-µm-pored membrane. Scale bar: 5.0 µm. **(d)** Ratio of mean fluorescence intensity (Vim/E-cad) in LifeAct-KCs located above the membrane or within the 3.0-µm micropores. For each micropore, fluorescence intensity was measured both above the membrane and within the pore, and the Vim/E-cad ratio was calculated. N = 10 micropores from three independent experiments. Data were analyzed with a paired t-test. ** p < 0.01. Representative images from three or more replicates per group are shown.

**Figure 3—figure supplement 1. LifeAct-KCs treated with chemicals**

**(a–c)** Representative images of LifeAct-KCs present within 3.0-µm micropores 6 hours after cell seeding. Cells were treated with TGF-β ligand **(a)**, TGF-β receptor inhibitors, SB431542, and SB525334 **(b)**, or cytochalasin D **(c)**. **(d–f)** Representative images of LifeAct-KCs present below 3.0-µm micropores 24 hours after cell seeding. Cells were treated with TGF-β ligand **(d)**, TGF-β receptor inhibitors **(e)**, or cytochalasin D **(f)**. Representative images from three or more replicates per group are shown.

**Figure 4—figure supplement 1. Genetic and pharmacological modulation of Piezo1 and keratin 6 in keratinocytes**

**(a, b)** Representative images of ruthenium red–treated LifeAct-KCs present within 3.0-µm micropores 6 hours after cell seeding **(a)** and below 3.0-µm micropores 24 hours after cell seeding **(b)**. **(c, d)** Representative images of LifeAct-KCs treated with Yoda1 present within 3.0-µm micropores 6 hours after cell seeding **(c)** and below 3.0-µm micropores 24 hours after cell seeding **(d)**. Scale bar: 30 µm. **(e)** qRT-PCR analysis of *PIEZO1* expression in parental (N = 9) and PIEZO1 KO (N = 7) KCs. Data were analyzed with two-tailed Mann–Whitney U tests. *** p < 0.001. **(f)** Western blot analysis using parental and PIEZO1 KO KCs. Immunoblotted by anti-PIEZO1 antibody and anti-GAPDH antibody. **(g)** qRT-PCR analysis of pan-*KRT6* expression in parental (N = 9) and pan-KRT6 KO (N = 7) KCs. Data were analyzed with two-tailed Mann–Whitney U tests. *** p < 0.001. **(h)** Western blot analysis using parental and pan-KRT6 KO KCs. Immunoblotted by anti-KRT6 antibody and anti-GAPDH antibody. **(i)** Representative images of parental LifeAct-KCs, PIEZO1 KO KCs, and pan-KRT6 KO KCs present within 3.0-µm micropores 6 hours after cell seeding (upper images) and below 3.0-µm micropores 24 hours after cell seeding (lower images). Scale bar: 30 µm. Representative images from three or more replicates per group are shown.

## Supplementary Movie Legends

**Supplementary Movie 1.**

A time-lapse movie of LifeAct-KCs that exited the 3.0-µm-pored membrane (related to Figure 2b). Images are each separated by 5 minutes.

**Supplementary Movie 2.**

A time-lapse movie of LifeAct-KCs that entered the 3.0-µm micropores (related to Figure 2d). Images are each separated by 5 minutes.

**Supplementary Movie 3.**

A time-lapse movie of LifeAct-KCs that entered the 3.0-µm micropores (high-magnified view; related to Figure 2e). Images are each separated by 1 minute.

**Supplementary Movie 4.**

A time-lapse movie of LifeAct-KCs that entered the 8.0-µm micropores (high-magnified view; related to Figure 2f). Images are each separated by 1 minute.

**Supplementary Table 1.**
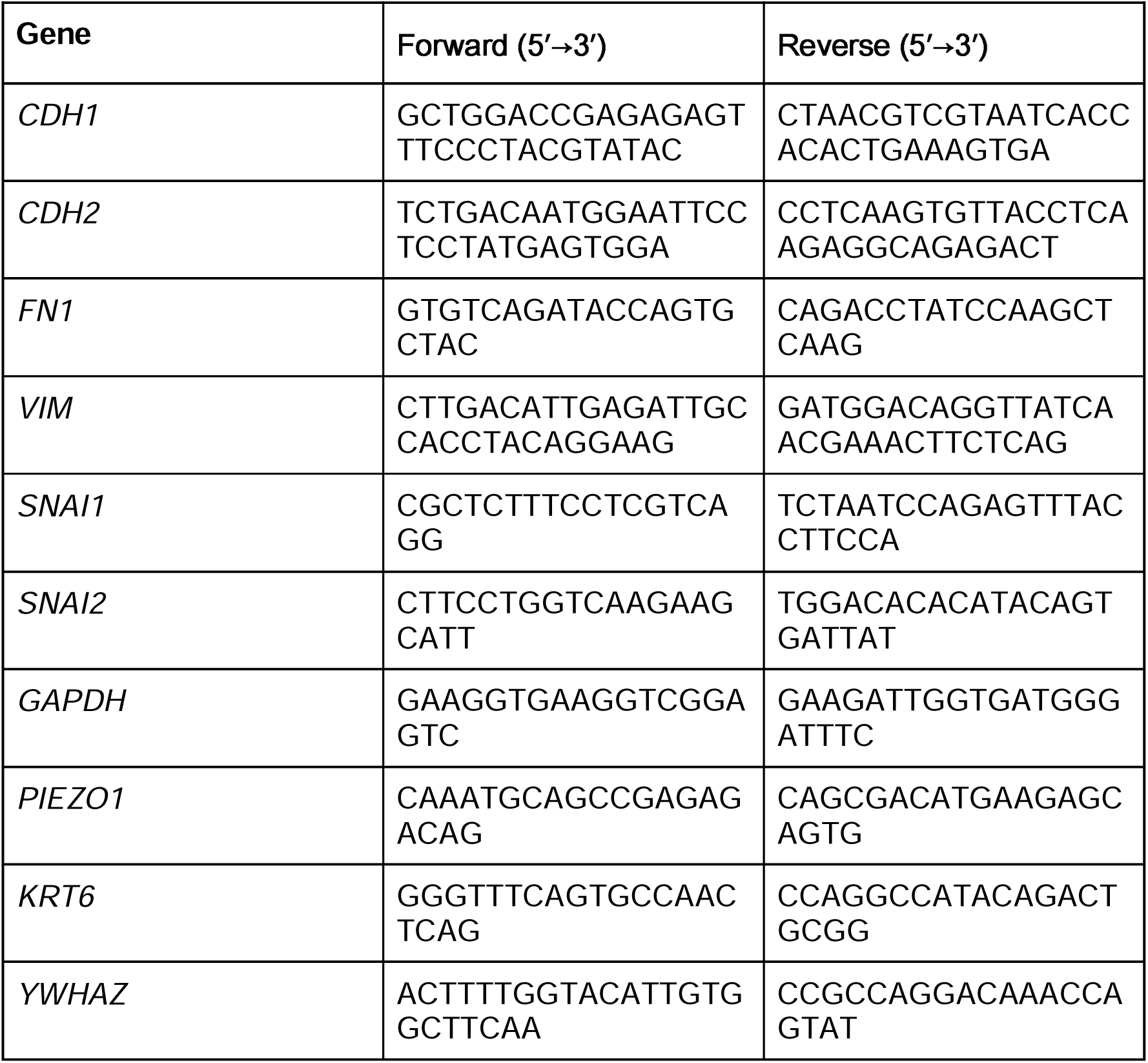
Primers used in qRT-PCR.

## Legends for Source Data

**Figure 4—source data 1**

Differentially expressed genes identified by bulk RNA-seq of HaCaT keratinocytes cultured on 0.4-µm- or 3.0-µm-pored membranes.

**Source data 1.** Numerical source data for all graphs.

Values plotted in Figure 1f, g; Figure 2h; Figure 3a–f; Figure 4a–f, l, m; Figure 1— figure supplement 1a–d; Figure 2—figure supplement 1d; and Figure 4—figure supplement 1e, g. Each worksheet is named after the corresponding panel. For the chemical treatment and knockout experiments, the experimental batch and the biological replicate are indicated for each value, and each value is the mean of eight measurement points, as described in Materials and Methods.

**Source data 2.** Uncropped images of the Western blots shown in Figure 4—figure supplement 1f and h.

Membranes with molecular weight markers for PIEZO1 and GAPDH (Figure 4— figure supplement 1f) and for KRT6 and GAPDH (Figure 4—figure supplement 1h). Lanes 1–3, parental cells; lanes 4–6, PIEZO1 KO or pan-KRT6 KO cells. M, molecular weight marker. Red dashed boxes indicate the regions shown in the figure.

