## Supplementary material for "Spatial confinement induces oscillatory migration of epidermal keratinocytes and forms a three-compartment epithelial structure with epithelial–mesenchymal transition dynamics": Figure 1-figure supplement 1

### Figure 1—figure supplement 1.
### Quantitative analysis of EMT marker expression and laminin β3 distribution

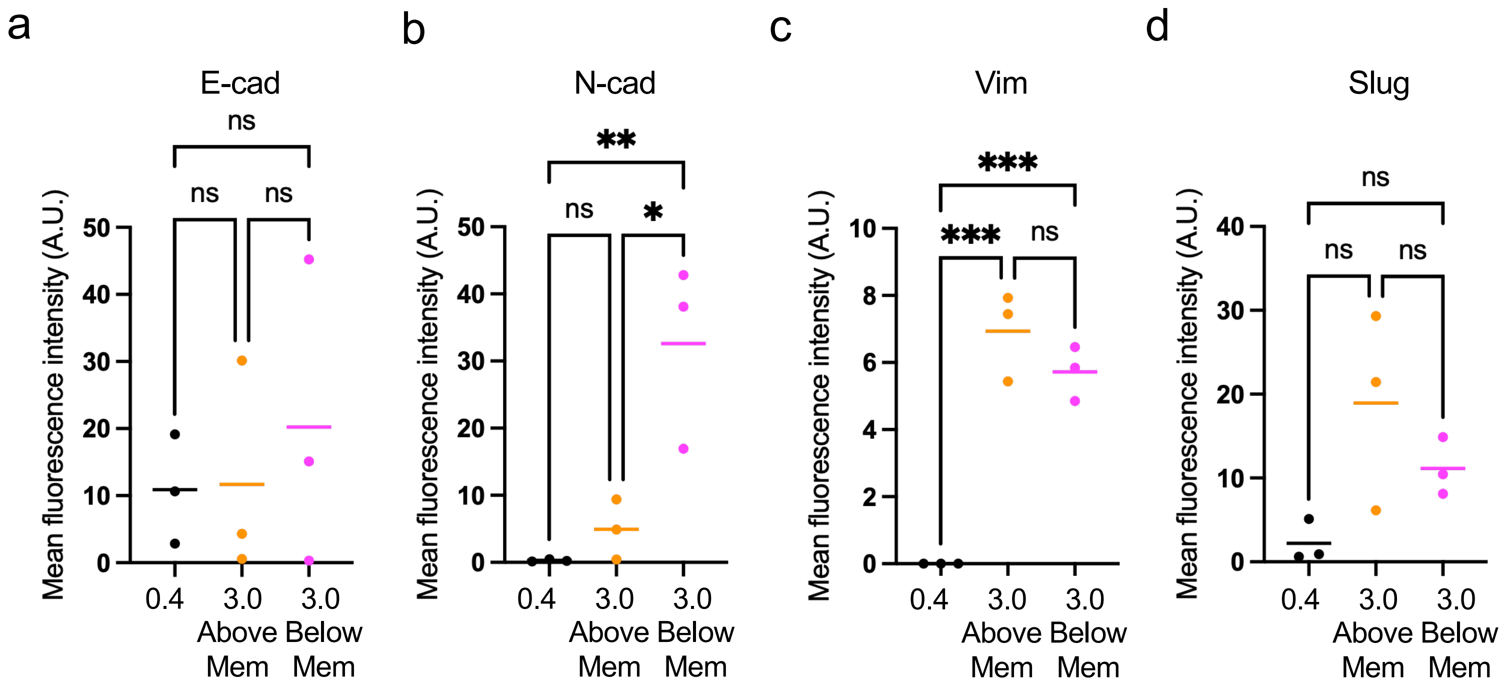


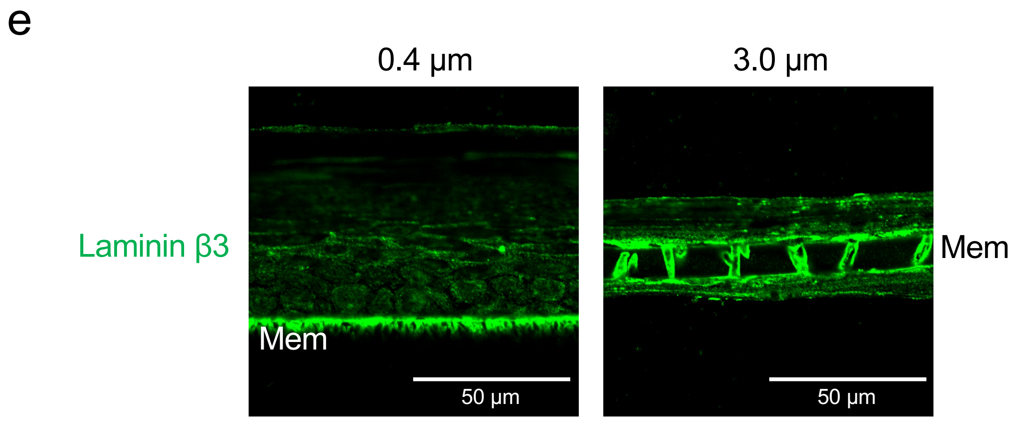
