## Supplementary figures and images for "Spatial confinement induces oscillatory migration of epidermal keratinocytes and forms a three-compartment epithelial structure with epithelial–mesenchymal transition dynamics"

### Figure 2-figure supplement 1

Figure 2—figure supplement 1.
Characterization of HaCaT KCs and LifeAct-KCs

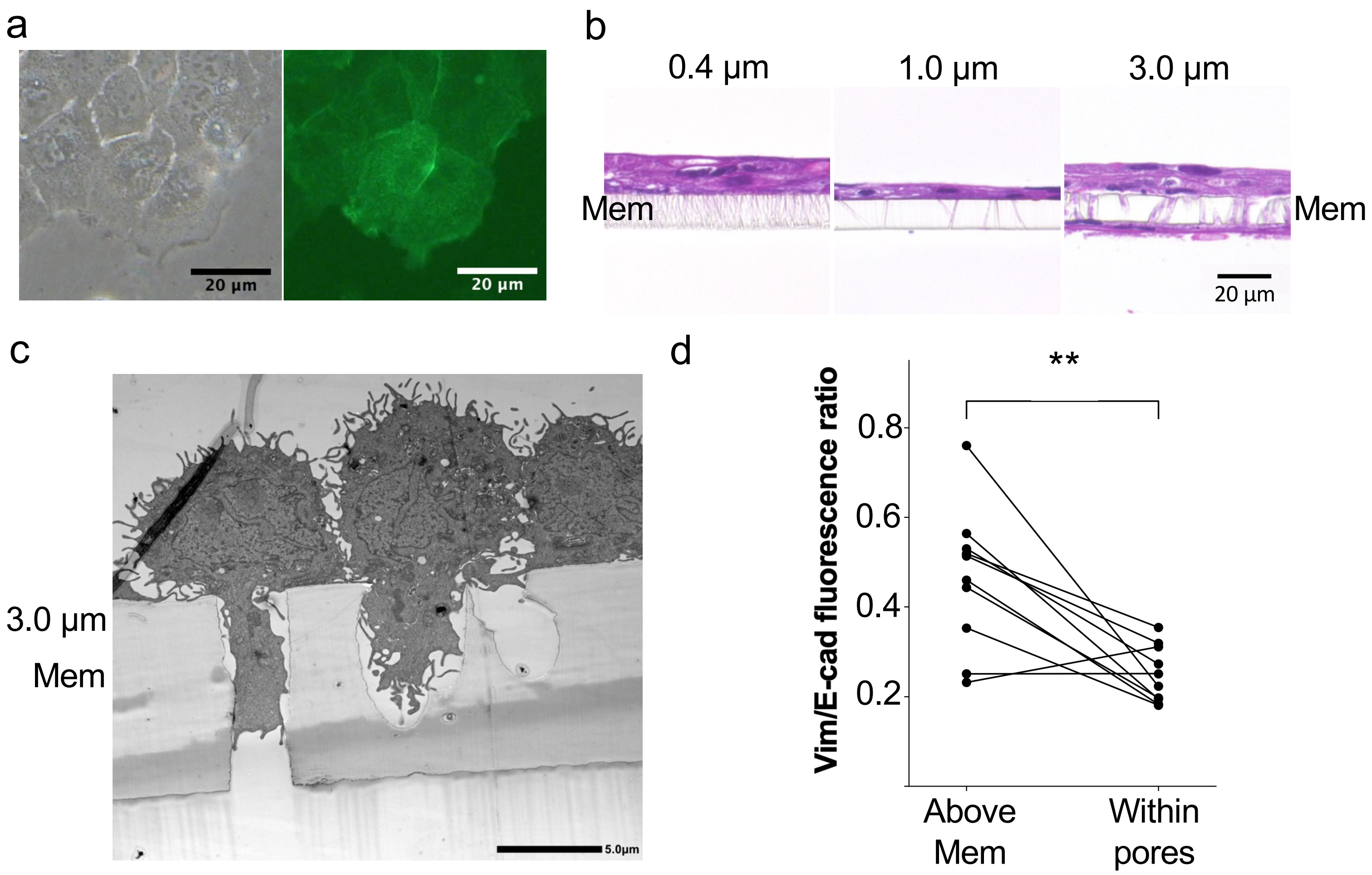

### Figure 3-igure supplement 1

#### Figure 3—figure supplement 1. LifeAct-KCs treated with chemicals

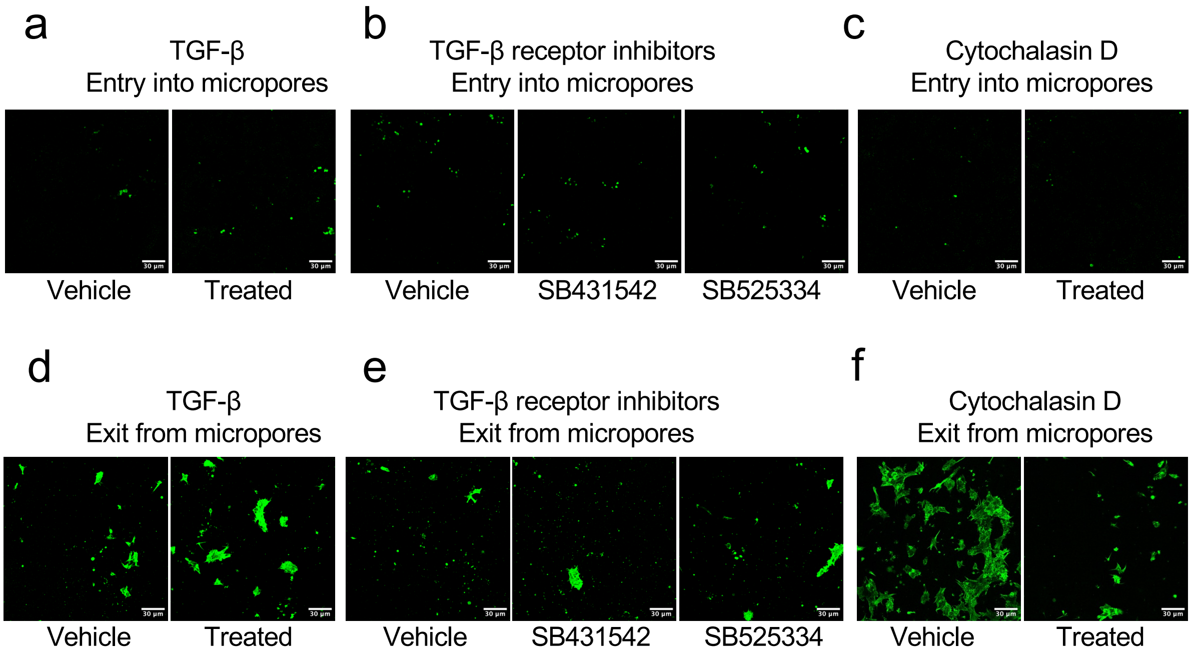

### Source data 2

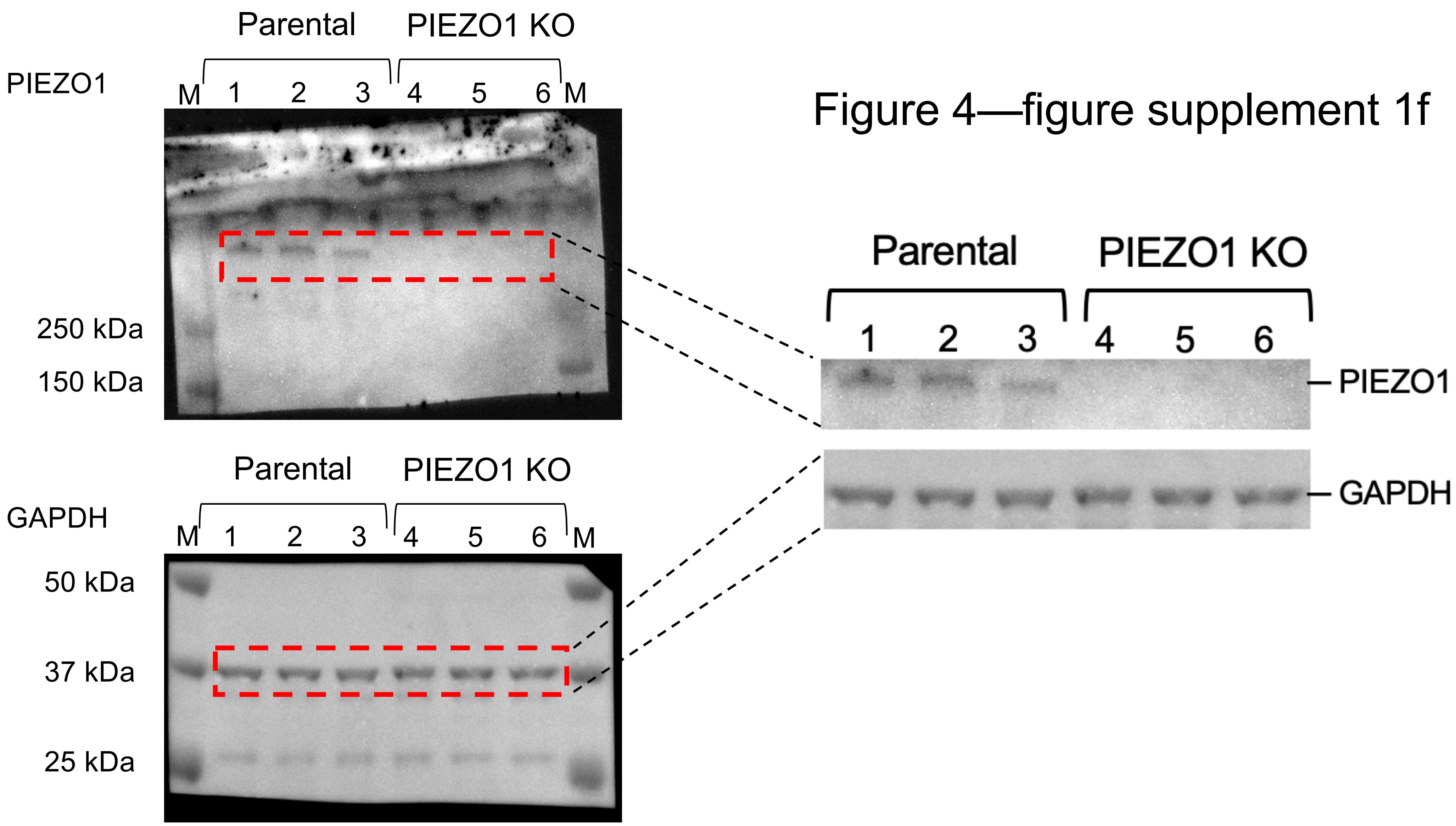

Figure 4—figure supplement 1h

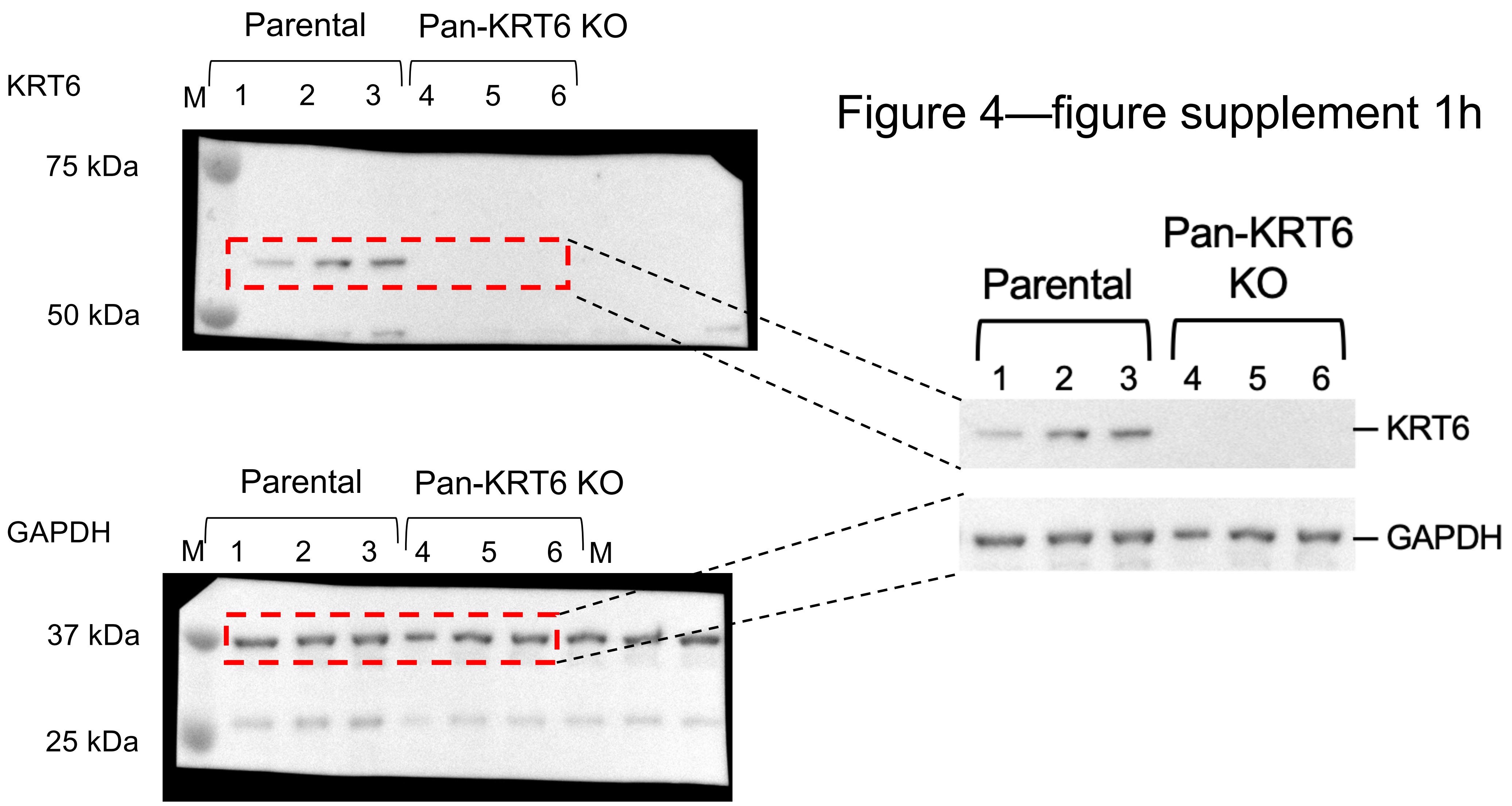
