## Supplementary material for "Spatial confinement induces oscillatory migration of epidermal keratinocytes and forms a three-compartment epithelial structure with epithelial–mesenchymal transition dynamics": Figure 4-figure supplement 1

### Figure 4—figure supplement 1. Genetic and pharmacological modulation of Piezo1 and keratin 6 in keratinocytes

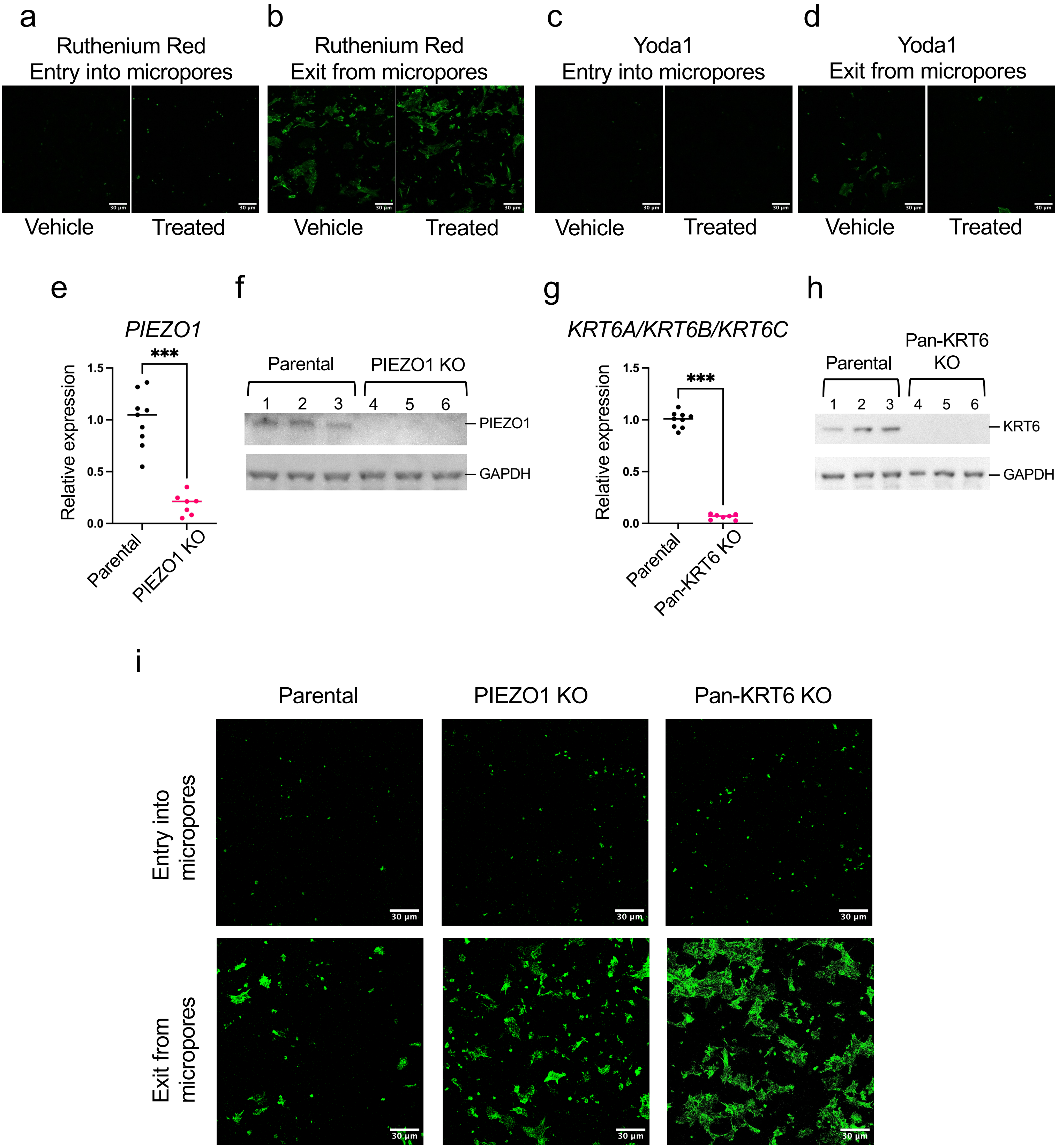
